# Autophagy at Crossroads: Modulating Responses to Combined Heat Stress and Bacterial Infection

**DOI:** 10.1101/2024.03.22.586360

**Authors:** Heike Seybold, Ella Katz, Yoram Soroka, Tamar Avin-Wittenberg

## Abstract

Plants face diverse stresses in natural environments, necessitating complex responses for survival. Abiotic and biotic stress responses are typically counteractive, posing challenges for breeding crops resilient to multiple stresses. Autophagy, a cellular transport process, plays a vital role in plant stress response, facilitating the degradation of cellular components and enabling nutrient recycling. Here, we asked what the role of autophagy is in combined abiotic (heat) and biotic (bacterial infection by *Xanthomonas campestris* pv. *vesicatoria*) stress. We introduce a conceptual framework based on assays monitoring autophagy activation, bacterial infection, and metabolic profiling.

We observed that heat stress facilitates bacterial growth in an autophagy-dependent manner. Bacterial effectors facilitate this phenomenon. We also demonstrate the engagement of the autophagy-related 8 (ATG8) protein family members in stress-specific activation. Metabolic profiling highlights effector-dependent shifts in nutrient availability during stress, influencing bacterial performance. Our study challenges the assumption that combined stresses are simply the sum of individual responses as exemplified by activation of the autophagic pathway. Instead, it establishes autophagy as a link connecting environmental factors and plant-microbe interactions. Insights for our study can present a novel perspective for designing strategies to enhance crop resilience in the face of multifaceted challenges.

## Introduction

Field-grown plants face many stresses, both abiotic (e.g., drought, temperature) and biotic (e.g., pathogens), requiring intricate physiological and molecular strategies for survival. Understanding these strategies has become more urgent due to climate change and the spread of new pests and diseases linked with these changes (Luck et al., 2011, Cavicchioli et al., 2019). Moreover, the nature and severity of stresses encountered by crops are predicted to worsen (Bebber, 2015, Hey et al., 2009). Abiotic and biotic stress responses are typically perceived to be counteractive, thus posing a significant challenge for breeding crops resilient to multiple stresses (Zhang and Sonnewald, 2017). Despite their distinct nature, there are intriguing similarities in the signaling patterns of biotic and abiotic stress responses, such as changes in phytohormone levels, calcium influx, reactive oxygen species (ROS) production, and activation of specific transcription factors (Atkinson and Urwin, 2012, Fraire-Velazquez et al., 2011, Jones and Dangl, 2006, Glazebrook, 2005, Dodds and Rathjen, 2010, Islam et al., 2018, Ohama et al., 2017, Zhu, 2016). Another common feature of these stresses is the redirection of energy resources from growth to stress responses (Signorelli et al., 2019).

Autophagy, a conserved eukaryotic mechanism, is vital for the recycling of cellular components and energy sourcing (Yang and Klionsky, 2009, Feng et al., 2013, Masclaux-Daubresse et al., 2020, Liu and Bassham, 2012). In plants, three types of autophagy have been described: Micro-, Macro-, and Megautophagy (Liu and Bassham, 2012, van Doorn and Papini, 2014, Marshall and Vierstra, 2018). Macroautophagy (hereafter called autophagy) is extensively studied and involves the formation of cytosolic double-membrane vesicles called autophagosomes. Autophagosomes deliver cytosolic cargo, which can be randomly (bulk autophagy) or selectively acquired (selective autophagy), to the vacuole for degradation (Liu and Bassham, 2012). Though autophagy functions in both plant biotic and abiotic stress responses (Signorelli et al., 2019), little is known regarding its role in combined stresses. Here, we focused on the role of autophagy in the combination of heat stress (abiotic stress) and bacterial infection (biotic stress).

Autophagy progression is divided into distinct steps and, in plants, involves more than 40 autophagy-related (ATG) proteins (Liu and Bassham, 2012, Yang and Klionsky, 2010, Chen and Klionsky, 2011, Chung, 2019, Wang et al., 2011). *ATG* genes are regulated on a transcriptional level, and *ATG* transcript availability and timing are critical for an appropriate stress response (Magen et al., 2022). Plant transcription factors were shown to confer abiotic stress resistance by inducing *ATG* gene expression and, consequently, autophagy (Wang et al., 2015, Zhu et al., 2018). The WRKY transcription factor family is particularly interesting regarding combined stress. WRKY33 induces autophagy in response to both abiotic and biotic stresses by activating the transcription of *ATG5* and *ATG7*. It is involved in heat tolerance in different plant species and, in addition to *ATG* genes, induces the expression of defense-related genes in *Arabidopsis thaliana* (Arabidopsis) (Lai et al., 2011, Schmidt et al., 2019, Mao et al., 2011, Lippok et al., 2007, Zhou et al., 2014, Zheng et al., 2006). Arabidopsis knockout mutants of *ATG* genes (*atg* mutants) are hypersensitive to different stresses, while overexpression of *ATG* genes improves plant fitness and performance (Liu and Bassham, 2012, Slavikova et al., 2008, Minina et al., 2018, Magen et al., 2022, Zhang et al., 2020).

Heat can have cytotoxic effects resulting in cellular metabolism and homeostasis perturbations due to protein denaturation and aggregation. Plant autophagy can remove heat-induced protein aggregates (Baxter et al., 2013, Zhou et al., 2013, Zhou et al., 2014). In addition, Neighbor Of BRCA1 (NBR1) mediates the selective degradation of proteins by autophagy to play a crucial role in the recovery from heat stress by regulating heat stress memory (Thirumalaikumar et al., 2020). NBR1 recruits cargo to the autophagosome by interaction with ATG8, a small ubiquitin-like protein, which undergoes lipidation by Phosphatidylethanolamine upon autophagy induction (Svenning et al., 2011, Zhou et al., 2013). The inner autophagosomal membrane, associated proteins (ATG8, NBR1), and cargo undergo degradation within the vacuole. The number of ATG8 isoforms varies among plant species, expanding from one isoform in algae to multiple isoforms in vascular plants (Zess et al., 2019). Notably, ATG8 exhibits a non-canonical function in response to heat stress, aiding the Golgi apparatus reassembly in Arabidopsis after acute heat stress (Zhou et al., 2023).

The role of autophagy in biotic stress response is more complicated and depends on the pathogen’s lifestyle (Sertsuvalkul et al., 2022). Bacterial plant pathogens, residing extracellularly, employ a type 3 secretion system (T3SS) to deliver effectors into the plant cell, manipulating cellular processes, including autophagy, for their benefit (Üstün et al., 2016, Lal et al., 2020, Leong et al., 2022b). NBR1-mediated selective autophagy can antagonize the effector-driven virulence (Leong et al., 2022b, Ustun et al., 2018). Besides bacterial pathogens, the oomycete *Phytophthora infestans* inhibits autophagy through effectors, countered by NBR1-mediated selective autophagy (Pandey et al., 2021, Dagdas et al., 2016, Dagdas et al., 2018). In addition, overexpression of *ATG5* or *ATG7* increased resistance against the necrotrophic fungal pathogen *Alternaria brassicicola*, and other examples support a positive role for autophagy in defense responses against necrotrophic plant pathogens (Minina et al., 2018, Lai et al., 2011, Lenz et al., 2011, Patel and Dinesh-Kumar, 2008). In line with these findings, autophagy plays a negative role in defense responses against obligate biotrophic pathogens (Wang et al., 2011). In contrast to oomycetes and bacterial pathogens, plant viruses, present intracellularly, are removed directly through NBR1-dependent selective autophagy (Hafren et al., 2017, Hafren et al., 2018).

Despite the significance of combined stresses in real-world natural and agricultural settings, research on their impact, particularly involving autophagy, remains limited. Understanding how plants respond to combined stresses is crucial for ensuring crop fitness and productivity in the face of climate change. The interaction between plant stress responses can be synergistic. Still, in most cases, responses to combined stresses differ significantly from those to single stresses (Rasmussen et al., 2013, Pandey et al., 2015, Ramegowda and Senthil-Kumar, 2015, Zhang and Sonnewald, 2017). Plant autophagy as a pathway involved in various stress responses has an exciting potential to increase plant tolerance towards combined stress scenarios with broad agricultural implications. Here, we investigate the pivotal role of autophagy in orchestrating the response to combined stresses. We chose a dual stress paradigm involving elevated temperature followed by infection with the bacterial pathogen *Xanthomonas campestris* pv. *vesicatoria* (*Xcv*), the causal agent of bacterial spot disease on pepper and tomato. Autophagy was shown to contribute to plant defense against *Xanthomonas* pathogens, while *Xcv* targets autophagy to inhibit appropriate defense responses (Yan et al., 2017, Leong et al., 2022b).

## Materials and Methods

### Biological material and growth conditions

*Nicotiana benthamiana* mutant line *Δroq1* was kindly provided by Johannes Stuttmann (CEA, France; (Grutzner et al., 2021)). Transient expression constructs were kindly provided by Yasin Dagdas (Gregor Mendel Institute, Austria; GFP-*StATG8* isoforms) and Elena Minina (Swedish University of Agricultural Sciences, Sweden; *AtATG7*). *Xanthomonas campestris* pv. *vesicatoria* (*Xcv*) strains 85-10 and 85-10 *ΔhrcV* were kindly provided by Guido Sessa (Tel Aviv University, Israel).

*Solanum lycopersicum* M82 and *N. benthamiana* were grown under a 16h light/8h darkness cycle (long day, LD), 22 °C. Tomato plants were used for stress experiments at the age of five weeks, tobacco plants were transformed and subsequently stressed at an age of approximately four weeks. All plant were grown in a mixture of commercial plant soil, perlit, and vermiculite (2:1:1).

### Stress treatments

To induce carbon starvation, plant trays were covered with aluminum foil for three days but kept in the same climate chamber as control plants without carbon starvation. For biotic stress treatments, *Xcv* suspensions of OD_600_ 0.2 in 10 mM MgCl_2_ were pressure-infiltrated into leaf tissue of *N. benthamiana* using a needle-less syringe. Plants were placed into a Percival chamber at 22 °C (control temperature) for 6 h. For heat stress treatments, plants were placed into a second Percival chamber and subjected to 40 °C for 6 h.

### Bacterial growth assay

*Xcv* was grown in NYGB medium at 28 °C overnight. For measuring bacterial growth, leaves of five-week old tomato plants or four-week old tobacco plants were pressure-infiltrated with *Xcv* OD_600_ 0.0004 in 10 mM MgCl_2_ with or without 5 mM 3-Methyladenine (3-MA) using a needle-less syringe. Leaf disk samples were taken immediately after infiltration of an independent set of plants to ensure equal infiltration densities. To check bacterial performance, leaf disk samples were taken three days after inoculation. Bacterial growth was monitored by serial dilution plating of ground leaf discs.

### Transient expression in tobacco

To study the stress specificity of ATG8 isoforms, *Agrobacterium tumefaciens* strain GV3101::pMP90 was transformed by electroporation with constructs of N-terminally GFP-tagged versions of all seven *St*ATG8 isoforms (Zess et al., 2019), respectively. For transient expression in *N. benthamiana Δroq1* plants, bacteria were grown in LB medium at 28 °C overnight, resuspended in infiltration buffer (10 mM MES, pH 5.7; 10 mM MgCl_2_; 150 μM Acetosyringone), adjusted to OD_600_ 0.5 and incubated for 2 hours at room temperature. The bacteria were subsequently pressure-infiltrated into *N. benthamiana* leaf tissue using a needle-less syringe. After an initial phase of 18 h under LD conditions for expression of constructs, the plants were subjected to the respective stress conditions.

### Protein extraction, SDS-PAGE, and Western blot

Equal amounts of plant leaf tissue were ground in liquid nitrogen using metal beads. Total proteins were extracted using a protein extraction buffer (50 mM Tris, pH 8; 20 mM NaCl; 10 % glycerol; 0.5 % Triton X-100; 1x Protease Inhibitor). Samples were mixed thoroughly, incubated on ice for 10 min, and centrifuged (14 000 rpm, 10 min, 4 °C). The supernatant was collected, mixed with SDS-PAGE sample buffer, and incubated at 80 °C for 5 min. Total proteins were separated by SDS-PAGE and transferred to a polyvinylidene difluoride membrane. Equal loading was validated using Coomassie Brilliant Blue staining of the membrane before the remaining unspecific protein binding sites were blocked with blocking solution (3 % milk + 2 % BSA in 1x PBST). For immunodetection, primary antibodies (anti-GFP: Abcam ab290) from rabbit were used in combination with a secondary HRP-conjugated anti-rabbit antibody from goat. For repeated detection of proteins from one membrane, membranes were first dried and then re-wetted with 1x PBST before Coomassie Brilliant Blue staining. The membranes were stripped by incubation in stripping buffer (200 mM glycine; 3.5 mM SDS; 1 % Tween 20; pH 2.2) for 20 min and subsequent washing with 1x PBST (3x for 5 min). Membranes were re-blocked with blocking solution prior to re-probing with the new antibody.

### GC-MS analysis

Plant metabolites from *N. benthamiana* were extracted following an established protocol for GC-MS-based metabolite profiling (Lisec et al., 2006), with minor modifications. In short, the samples (25 mg leaf tissue per sample with six biological replicates per treatment) were extracted in methanol containing an internal standard (Pentaerythritol, stock concentration 0.6 mg/ml in DDW, final concentration 0.016 mg/ml), shaken for 15 min at 70 °C, and centrifuged at 14 000 rpm for 10 min. The supernatant was transferred to new tubes, and chloroform and double-distilled water were added. The mixture was centrifuged at 14 000 rpm for 15 min, and aliquots of 200 μL of the upper phase of each sample were transferred to new tubes and dried by Speedvac, overnight. Derivatization was carried out as previously described (Lisec et al., 2006). Polar metabolites were measured by the Agilent 7200B GC/Q-TOF. The injection and separation procedures were performed according to Dahtt using the DB-35MS column (Dhatt et al., 2019). Metabolite detection and annotation were performed by Quant software (Agilent, Santa Clara, CA, USA), according to an accurate mass library of known markers generated by our group and run in the same system. Following blank subtraction, the peak area of each metabolite was normalized to the internal standard (i.e., Pentaerythritol) in each sample, as well as to the fresh weight of the respective sample. For more information, see Table S1.

### Statistical analysis and data visualization

Data were statistically examined using analysis of variance (Ordinary one-way ANOVA) and tested for significant differences using unpaired *t* tests using GraphPad Prism 8.0.1 software. Graphs and were generated using GraphPad Prism 8.0.1, while Principal component analysis (PCA) of the metabolome was performed using Metaboanalyst, an open-source, web-based tool for metabolomics data analysis (https://www.metaboanalyst.ca/). Heatmaps were generated using the MultiExperiment Viewer (MeV) freely available software application (Howe at al, 2010). Western blots were processed to maximize readers accessibility to the data. Original blots can be found in the supplemental data (FigS3).

## Results

### Heat stress facilitates bacterial growth in an autophagy-dependent manner

Autophagy functions in plant biotic and abiotic stress response (Avin-Wittenberg, 2019, Sertsuvalkul et al., 2022). Yet, very few studies investigated the role of autophagy in a combined stress scenario, which is closer to the conditions faced by plants in the field. We chose to examine two stresses in which autophagy was shown to play a role: heat stress and *Xcv* infection (Leong et al., 2022b, Zhou et al., 2014). We began with assessing the growth of *Xcv* following infiltrated with *Xcv* following 6h heat stress. Bacterial titers were checked 3 days post-infection (dpi) (Fig1A). *Xcv* grew better in heat-stressed plants (Fig1B). Surprisingly, this beneficial effect of preceding heat stress on bacterial growth was dependent on autophagy. In the presence of the autophagy inhibitor 3-Methyadenine (3-MA), preceding heat stress did not affect bacterial growth. Interestingly, pre-treatment with 3-MA without heat stress did not alter bacterial growth. We thus postulated that activation of autophagy during heat stress facilitates bacterial growth.

**Figure 1:**
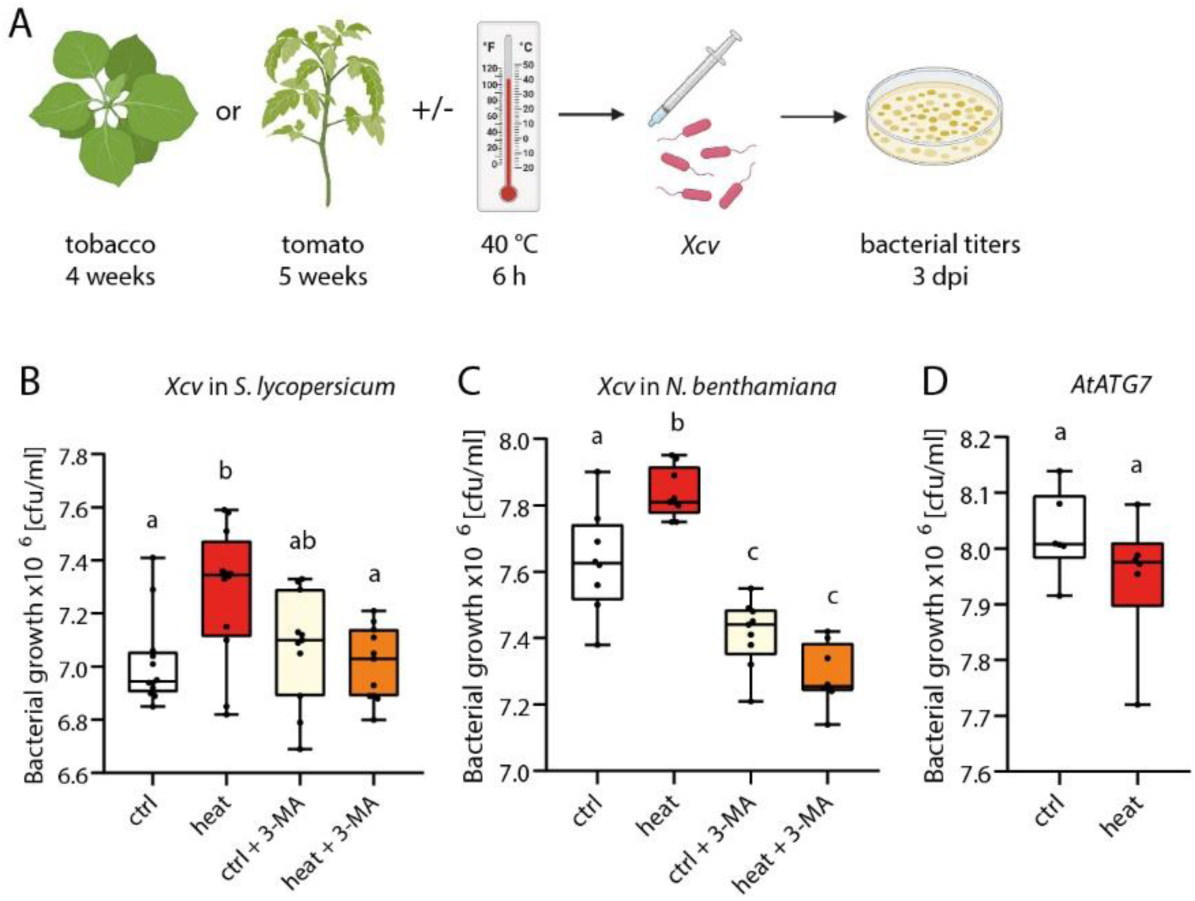
*Xcv* growth is benefited by preceding heat stress in an autophagy-dependent manner. (**A**) Schematic representation of the experimental setup to test bacterial growth in previously heat-stressed plants. (**B,C**) Bacterial growth was tested in tomato (**B)** or tobacco (**C**) plants that were previously subjected to heat or control temperature and co-infiltrated with the autophagy inhibitor 3-MA. (**D**) Bacterial growth in heat- and control-treated tobacco plants transiently expressing *AtATG7*. Experiments were carried out independently at least twice (B)/ three times (C, D) with similar results. Number of biologically independent replicates: *Xcv* in tomato (*n* = 12), *Xcv* in tobacco (*n* = 9), *At*ATG7 (*n* = 6). Statistical analysis was performed by One-way ANOVA and Turkey’s multiple comparison test.

Over-expression of ATG7 was shown to increase autophagic activity in Arabidopsis (Minina et al., 2018). Thus, we aimed to overexpress ATG7 and assess bacterial growth. To that end, we switched to a *Nicotiana benthamiana* system, in which autophagy following *Xcv* infection has been previously investigated and transient overexpression is easily performed (Leong et al., 2022b). Although *N. benthamiana* is not a natural host of *Xcv*, the mutant line *Δroq1* (Grutzner et al., 2021) can be infected by the bacteria. We initially examined whether the phenomenon observed in tomato also occurs in *N. benthamiana*. We infiltrated *Xcv* into *N. benthamiana Δroq1* plants after 6h heat stress. As in the tomato system, we observed increased bacterial growth following heat stress, which was attenuated by autophagy inhibition (Fig1C). The differences in the *N. benthamiana* system were even more pronounced than in tomato. We observed a slight yet significant decrease in bacterial growth following 3-MA treatment under control conditions (Fig1C). We thus wished to verify that the 3-MA treatment itself did not affect bacterial growth. We incubated *Xcv* in an infiltration medium with and without 3-MA and pated the bacteria directly to assess growth. There was no difference in bacterial titters between treatments (control: log10cfu/cm^2^ 4.25±0.11, 3-MA: log10 cfu/cm^2^ 4.28±0.08, n=4, no significant difference found in student’s *t-test*). Next, we tested bacterial performance with and without preceding heat stress in *N. benthamiana* plants transiently expressing *AtATG7*. As expected, in plants transiently overexpressing *AtATG7*, *Xcv* grew similarly in heat-stressed and unstressed plants (Fig1D), suggesting that autophagy activation, irrespective of its cause, is conducive to bacterial growth.

### Different ATG8 homologs are involved in stress-related autophagy

As we postulated that autophagy activation by heat stress before *Xcv* infection benefits bacterial growth, we wished to determine that autophagy is indeed activated under these conditions. We focused on ATG8, which is anchored to the autophagosome membrane. The GFP-ATG8 fusion construct is used as a common autophagosome marker. Moreover, as ATG8 is degraded in the vacuole, the ratio of GFP-ATG8 and free GFP resealed following degradation is used as a proxy of autophagic flux (Klionsky et al., 2021). ATG8 exists as a gene family in plants and also functions in target recognition and cargo binding for selective autophagy. We speculated that ATG8 isoform specificity may vary between stresses, which is yet to be determined (Kellner et al., 2017). We thus examined the degradation patterns of GFP-fusion constructs of the potato (*Solanum tuberosum*) *St*ATG8 isoforms under single and combined stresses. As our chosen biotic stress is a tomato pathogen, potato isoforms were used to mimic the tomato response to *Xcv* infection.

GFP-tagged versions of all seven *St*ATG8 isoforms were transiently expressed in *N. benthamiana*. The plants were then subjected to heat stress, *Xcv* infection, and prolonged darkness. Prolonged darkness leads to carbon starvation, widely known to induce autophagy, presumably in a non-selective manner (Barros et al., 2017), and was therefore used as a positive control. Carbon starvation was induced by three days of darkness, leading to increased degradation of all GFP-*St*ATG8 isoforms and high levels of free GFP (Fig2A). Heat stress reduced protein levels of all GFP-*St*ATG8 isoforms after 6 h and 8 h of 40°C, yet we were not able to observe a free GFP band in all the samples, and thus compared Protein levels to the non-treated (control) plants (Fig2B). However, ATG8 degradation differed in timing and intensity. While some isoforms responded quickly (after 6 h), others showed a slower reduction upon stress (after 8 h). GFP-*St*ATG8-3.1 and GFP-*St*ATG8-4 showed a quick, strong, and lasting decrease in protein levels.

**Figure 2:**
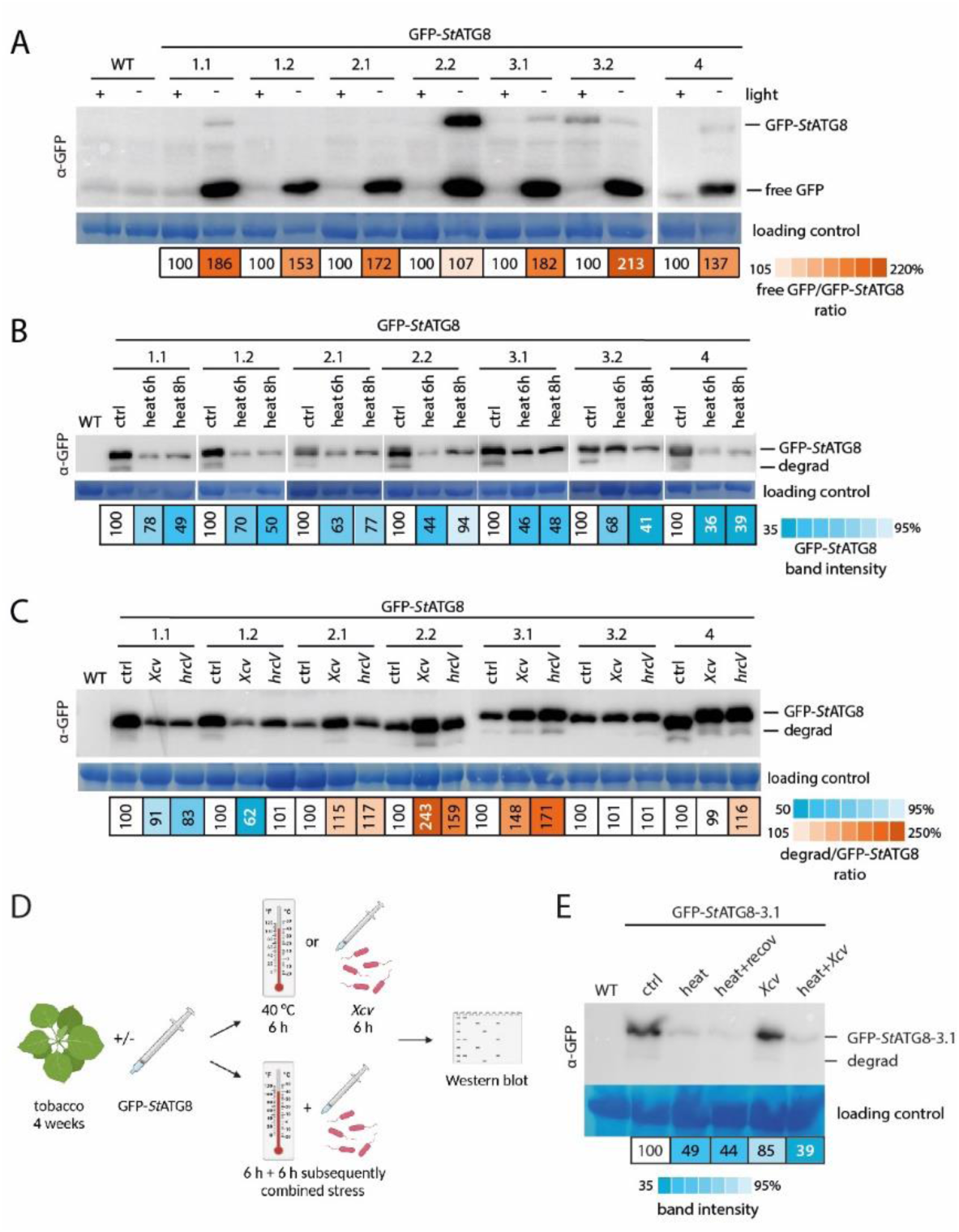
Autophagy is differentially activated under combined stress. (**A-C)** All seven GFP-*St*ATG8 isoforms were transiently expressed in *N. benthamiana*. The plants were subjected to different stresses 1 day after syringe-infiltration of GFP-*St*ATG8 constructs. Western blots show the degradation pattern after 3 days of darkness (**A**), 6/8 h of 40 °C (**B**) and 6 h after *Xcv/hrcV* infection (**C**). Degradation was quantified via measurement of band intensities of GFP-*St*ATG8 (**B**) or GFP-*St*ATG8 and free GFP (**A**) or a specific degradation band (**C**), respectively. (**D**) Schematic representation of the experimental setup to test the effect of combined heat-*Xcv* stress of GFP-*St*ATG8-3.1 degradation and gene expression. (**E**) Degradation pattern of transiently expressed GFP-*St*ATG8-3.1 upon heat, *Xcv,* and a combined heat-*Xcv* challenge. Band intensities were normalized to the protein loading control, and ratios of GFP-*St*ATG8 and a degradation band (if applicable) were calculated. Original blots can be found in the supplemental Fig S3. The experiments were carried out independently three times (B, C, E)/twice (A) with similar results.

Next, we tested the effect of infection with *Xcv*. Only a subset of GFP-*St*ATG8 isoforms was degraded upon *Xcv* challenge (Fig2C). The GFP-*St*ATG8-2 isoforms, GFP-*St*ATG8-3.1, and in some but not all experimental replicates, GFP-*St*ATG8-4 displayed increased degradation upon *Xcv* infection. GFP-*St*ATG8-1 isoforms showed the opposite effect. Interestingly, the *Xcv*-specific activation of GFP-*St*ATG8 isoforms was independent of bacterial effectors. Infection with the T3SS mutant *Xcv ΔhrcV* (hereafter called *hrcV*), lacking the effector secretion system (Cerutti et al., 2017), resulted in a GFP-ATG8 degradation pattern similar to that of *Xcv*. Surprisingly, the detected degradation band was not free GFP, unlike the observed for carbon starvation. These results indicate a specific autophagic response to stress, particularly to *Xcv*/bacterial infection, unlike the non-specific reaction to general stressors like carbon starvation.

### Differential autophagy activation under combined stress compared to single stresses

As *St*ATG8-3.1 was degraded upon heat stress as well as after inoculation with *Xcv*, we chose this isoform to assess autophagic activity in a combined heat-*Xcv* challenge. As performed in the single-stress experiments, GFP-*St*ATG8-3.1 was transiently expressed in *N. benthamiana Δroq1* plants. The plants were then subjected to 6 h heat stress or 6 h *Xcv* infection. In addition, 6 h heat stress was followed by subsequent 6 h *Xcv* or a 6 h recovery period without *Xcv* inoculation (Fig2D). Although *Xcv* inoculation was the second and, therefore, more recent treatment, GFP-*St*ATG8-3.1 degradation after combined stress treatment resembled heat stress instead of *Xcv* infection (Fig2E). Our results suggest that different ATG8 isoforms participate in autophagy activation during different stresses and that stress combinations can further influence autophagic flux in a non-additive manner. In the case of heat-*Xcv* challenge, preceding heat indeed changed the nature of autophagy activation during infection, further strengthening our hypothesis that pre-activation of autophagy promotes *Xcv* proliferation.

### Heat stress recovery fails to halt bacterial growth, but effector deficiency succeeds

When we examined autophagy activation upon combined stresses, we also noticed that the degradation pattern of GFP-ATG8 remained similar to heat stress after a 6h recovery period (Fig2E). We thus wished to assess whether the phenomenon of increased bacterial growth following heat stress also persisted after recovery. We, therefore, repeated bacterial growth experiments with heat-stressed plants and prolonged the recovery period to 24 h (Fig3A). Comparable to results without a recovery period, previous heat treatment resulted in increased bacterial growth even after the recovery period (unpaired *t-test* p-value 0.0067), although not significantly different in the ANOVA (Fig3B). As seen without the recovery period, this effect was again autophagy-dependent. These results suggest that autophagy induction upon heat stress has a lasting effect on the internal plant environment, impacting bacterial growth.

**Figure 3:**
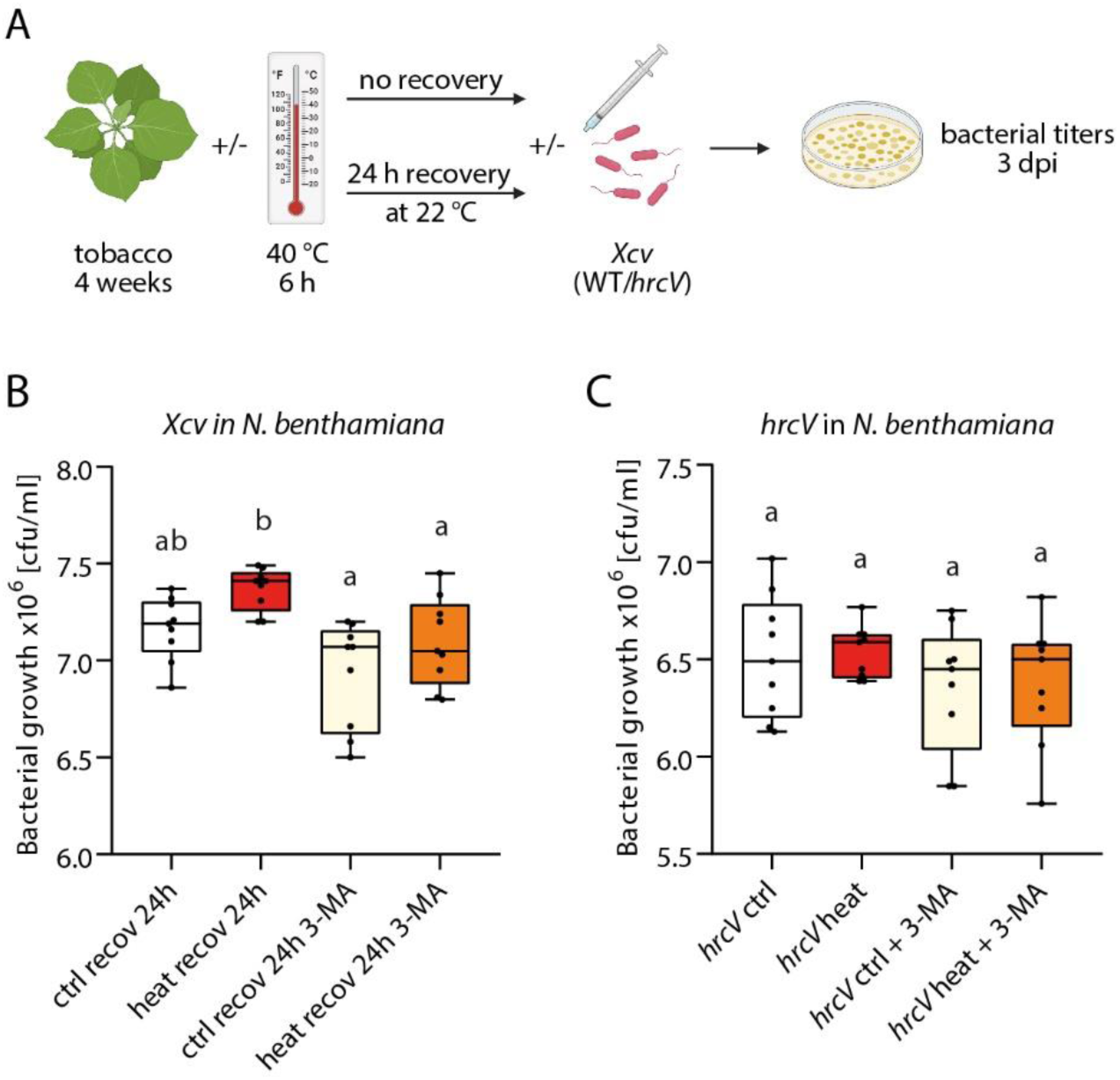
Increased bacterial growth after heat stress is not diminished by a recovery period but is effector-dependent. (**A**) Schematic representation of the experimental setup to test the effect of a recovery phase or effector dependency on bacterial growth. (**B,C**) Bacterial growth of *Xcv* (**B)** or *hrcV* (**C)** was tested in tobacco plants. (**B**) plants were previously subjected to heat or control temperature followed by a 24 h recovery period at control temperature. (**C**) plants were subjected to heat or control conditions followed immediately by bacterial infiltration. In both cases, plants were co-infiltrated with the autophagy inhibitor 3-MA (*n* = 9 in both experiments). Statistical analysis was performed by One-way ANOVA and Turkey’s multiple comparison test.

Our Western blot analyses also revealed that autophagy activation by *Xcv* is effector-independent (Fig2C). Thus, we wished to examine whether effectors play a role in autophagy-mediated bacterial growth and examined the *hrcV* mutant following heat stress. Interestingly, the mutant did not benefit from previous heat stress in the host plants (Fig3C), and autophagy inhibition by 3-MA did not affect its growth either. We thus postulate that although autophagy is activated by bacterial infection in an effector-independent manner, the autophagy-dependent increase in bacterial growth after heat stress is also dependent on bacterial effector delivery.

### Metabolic shift caused by *Xcv* is treatment-resilient

Recognizing autophagy’s role in plant nutrient availability (Signorelli et al., 2019), we theorized that its activation during heat stress might reshape metabolic content to favor bacterial growth. We performed metabolic profiling via Gas Chromatography-Mass Spectrometry (GC-MS) on concentrated on primary metabolites, representing nutrients potentially utilized by the bacteria. Principal component analysis (PCA) showed a major shift between non-treated plants (red) and samples from heat-treated plants (yellow) (Fig4B). *Xcv*-inoculation led to another shift. All samples that received *Xcv*-treatment clustered together independent of additional treatments. Samples from *hrcV*-infected plants were also analyzed to assess the influence of effectors on metabolism. Most of the *hrcV*-treated samples clustered with the *Xcv*-treated samples, yet some clustered away from the *Xcv*-treated samples in the PCA, pointing to a slight effector-dependent shift of the metabolome (Fig4B). We also included samples in which plants had a 24 h recovery period from heat before infection. In accordance with the bacterial growth assay, the recovery period did not affect the metabolic profile of the samples (FigS1).

**Figure 4:**
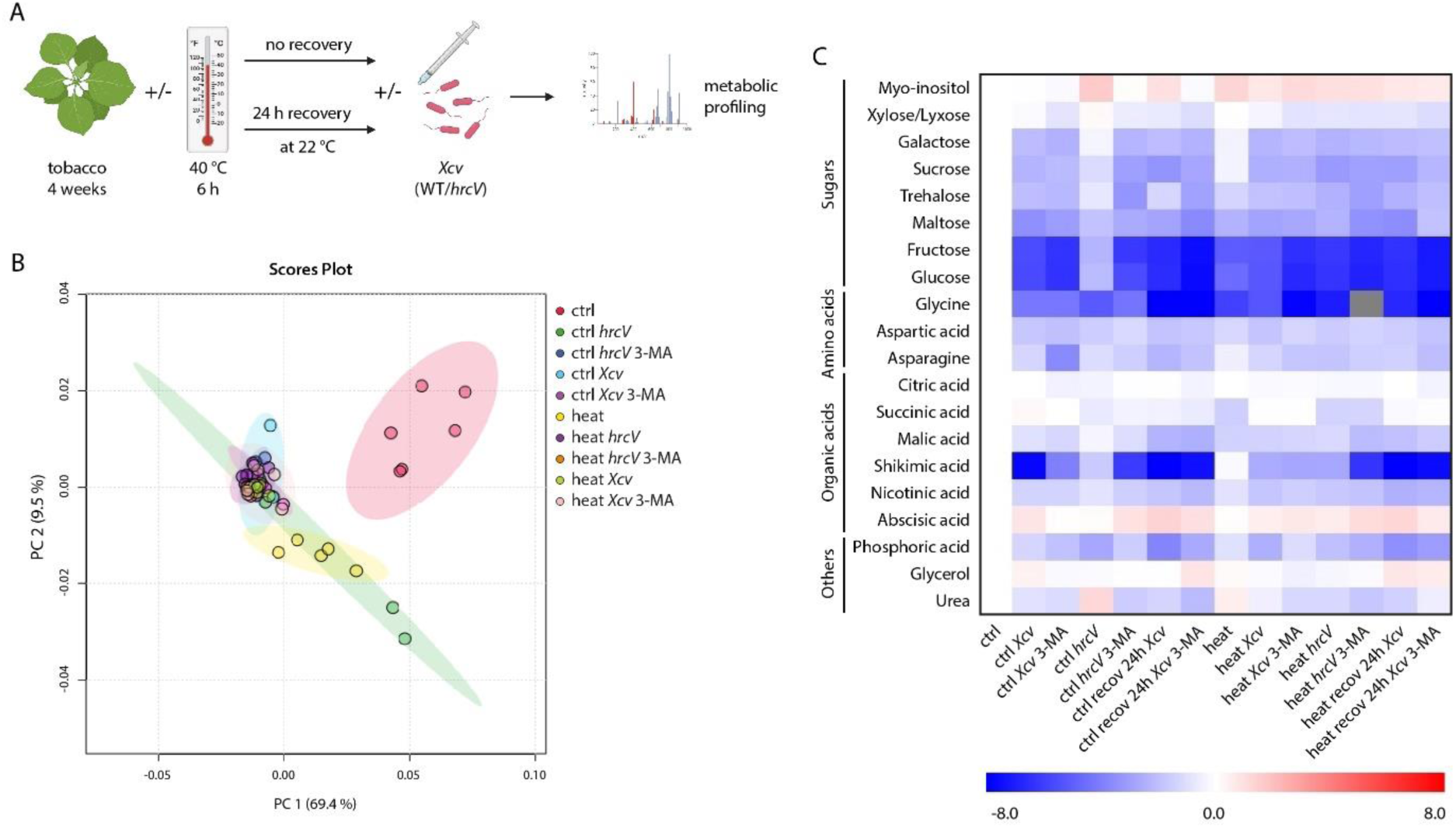
Bacterial infection causes pronounced effects on the plant metabolome. (**A**) Schematic representation of the experimental setup conducted for metabolic profiling. *N. Benthamiana* plants were exposed to heat stress with/without a 24 h recovery period, then infiltrated by *Xcv/hrcV* and co-infiltrated with the autophagy inhibitor 3-MA. Samples were collected for metabolic profiling by GC-MS 3 d post-infection (n=5-6). (**B)** Principal component analysis of the metabolome in response to heat treatment or bacterial infection with and without a recovery phase and 3-MA treatment. (**C**) Heat maps displaying median values in log2 scale of the response of 4 different groups of metabolites to stress treatment as in (A). Red color refers to upregulation compared to the control; blue color refers to downregulation compared to the control.

Generally, heat-treated plants as well as *Xcv*-infected plants, showed strong differences from control plants in most identified compounds (Fig4C). Many of the metabolites were depleted. In particular, sugars and amino acids were reduced under heat stress and after infection, respectively. In contrast, the sugar alcohol Myo-inositol increased, specifically after heat challenge (see below). Collectively, our findings indicate stress-induced metabolic shifts, with stress-specific patterns yet common alterations in energy sources. Notably, the response to *Xcv* infection exhibited remarkable resilience to additional factors or stresses.

### Changes in organic acids are independent of autophagy

We next focused on specific metabolic changes induced by stress combinations and their dependence on autophagy. No changes in shikimic acid levels were detected in response to heat stress, but levels were reduced significantly in response to infection or a combined heat-pathogen stress (FigS2A). This response to infection was independent of autophagy and similar in *Xcv* and *hrcV*. Nicotinic acid showed a mild reduction under combined stress compared to the single stresses, which was significantly lower than in control plants (FigS2B). Again, the response to combined stress was independent of autophagy. Two compounds of the TCA cycle, succinic acid and malic acid, were reduced only upon heat treatment but not infection (FigS2E,F). However, while levels of malic acid stayed low in all plants that previously received heat treatment, *Xcv* but not *hrcV* infection restored control levels of succinic acid in heat-stressed plants in an autophagy-independent manner. The levels of a third detected member of the TCA cycle, citric acid, did not change in response to any stress treatment (FigS2G). Together, our results suggest that changes in organic acids in response to stress combinations are not dependent on autophagy.

### Shift in amino acid levels is independent of bacterial effectors

We directed our attention to metabolites functioning as energy sources. Amino acids are a major energy source that can be mobilized under stress (Signorelli et al., 2019). We detected three amino acids, which were generally reduced upon bacterial infection as well as heat stress, especially in the case of glycine (Fig5A,B). *Xcv* and *hrcV* showed a similar effect. Asparagine was reduced upon bacterial infection and further reduced in the presence of 3-MA under control conditions (Fig6C). This effect was also observed in heat-stressed plants but to a lesser extent. Thus, the shifts in amino acid levels are not a result of bacterial manipulation but rather a general response of the plant to stress conditions. Urea levels, a metabolite directly linked to amino acid catabolism (Witte, 2011), stayed stable upon heat challenge but were reduced upon *Xcv* infection, independent of temperature (Fig6D). Interestingly, *hrcV* treatment significantly increased urea levels in an autophagy-dependent and heat-sensitive manner.

**Figure 5:**
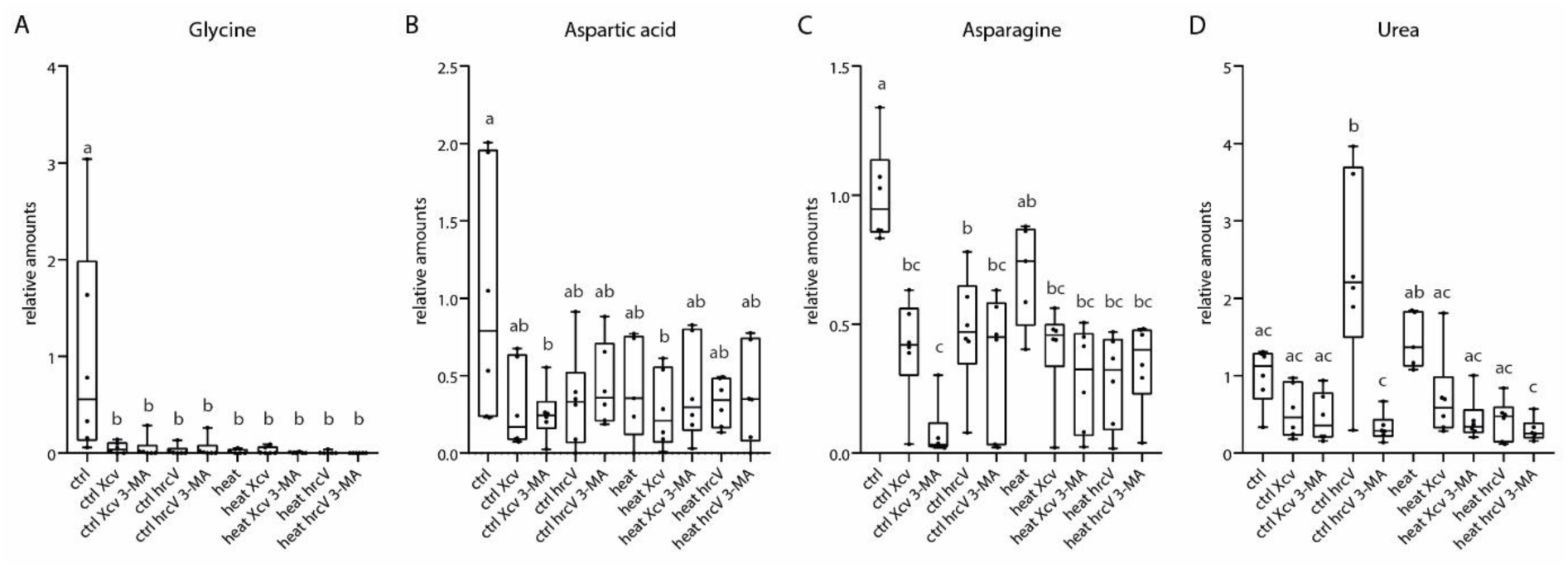
Detailed overview of effects of *Xcv* infection on the levels of three amino acids and urea. Boxplots showing levels of Glycine (**A**), Aspartic acid (**B**), Asparagine (**C**), and urea (**D**) in response to *Xcv*, *hrcV*, and heat treatment, respectively (*n* = 5-6). Statistical analysis was performed by One-way ANOVA and Turkey’s multiple comparison test.

**Figure 6:**
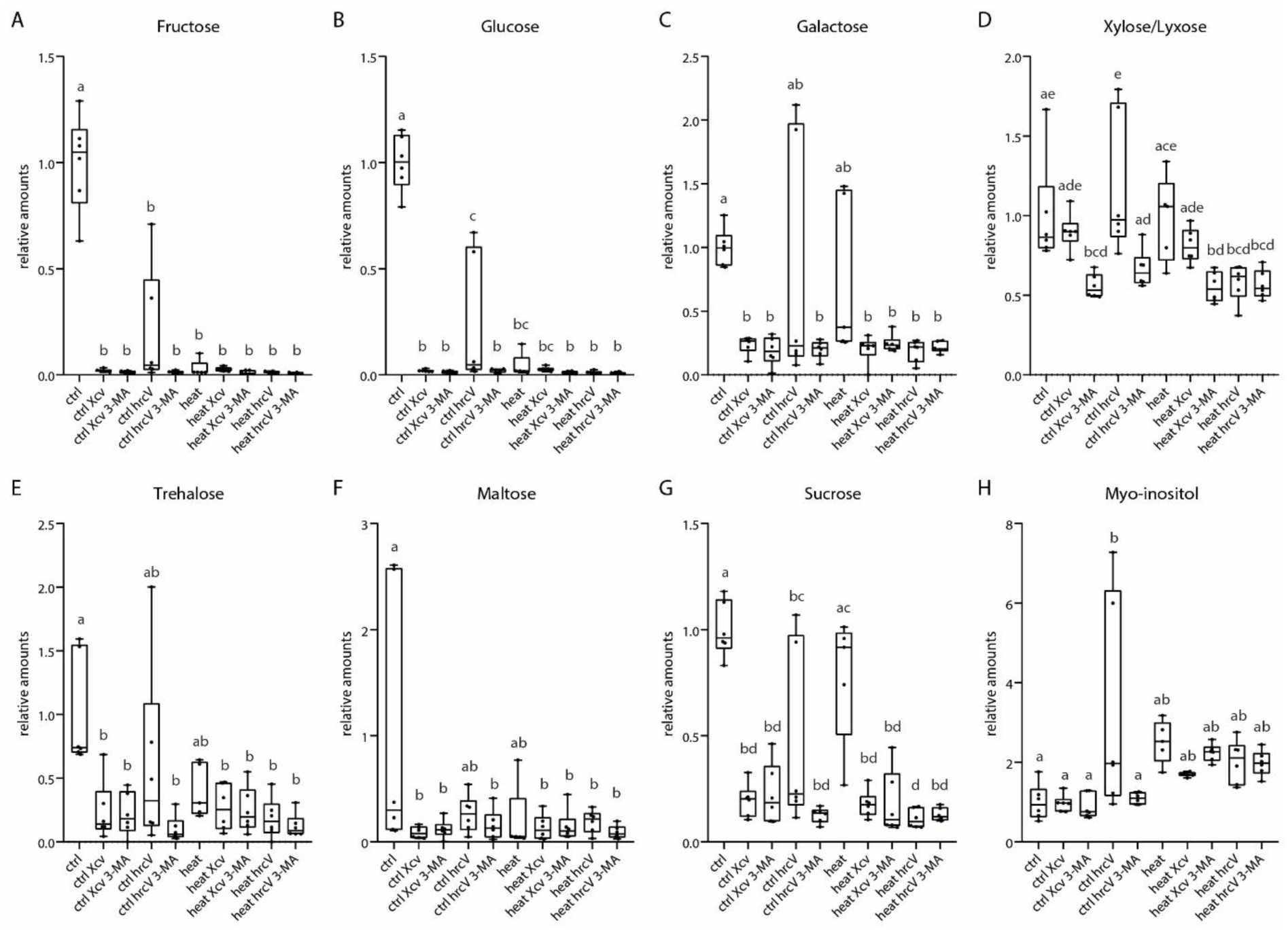
Detailed overview of effector-dependent effects of *Xcv* infection on sugar levels. Boxplots showing levels of (**A-D**) Mono-, (**E-G**) Disaccharides, and (**H**) sugar alcohol in response to *Xcv*, *hrcV*, and heat treatment, respectively (*n* = 5-6). Statistical analysis was performed by One-way ANOVA and Turkey’s multiple comparison test.

### Effector-dependent reduction of sugars is heat-sensitive and modulated by autophagy

Sugars are a main energy source under stress conditions (Signorelli et al., 2019). Most of the mono- and disaccharides were globally decreasing upon heat- and *Xcv*-treatment compared to control samples, with sucrose showing only a decrease upon infection, not heat (Fig7A-C, E-G). This effect was not influenced by 3-MA treatment. Samples treated with *hrcV* displayed a more minor sugar reduction than *Xcv* infection (significant only for Glucose). However, when autophagy was inhibited by 3-MA treatment, *hrcV* infection reduced sugar levels to a similar extent as after *Xcv* infection. The effect of *Xcv* treatment was not further intensified by 3-MA treatment, as seen for *hrcV*. We, therefore, suggest that the decrease in sugar levels upon infection is partially dependent on effector(s) targeting autophagy. Interestingly, the effect observed for *hrcV* was only present in control conditions and absent in heat-stressed plants where *Xcv* and *hrcV* infection had similar, autophagy-independent effects.

The only detected Monosaccharide that did not fit into the described pattern is Xylose/Lyxose (Fig7D). Levels of this sugar were not reduced upon *Xcv* infection or heat treatment but only upon inhibition of autophagy, pointing to a role of autophagy in maintaining Xylose/Lyxose levels under stress conditions.

Myo-inositol did not change upon *Xcv* infection (Fig7H). However, levels increased significantly upon *hrcV*-treatment in an autophagy-dependent manner. Heat stress led to a general increase in Myo-inositol levels, a well-documented response to heat and other abiotic stresses in different plant species (Khurana et al., 2017, Sharma et al., 2020, Boominathan et al., 2004). Combined heat-*Xcv* but not heat-*hrcV* stress reduced these levels significantly in an autophagy-dependent manner (ANOVA performed in the absence of control *hrcV* samples). The massive shifting of sugars is, at least partially, the result of an effector-mediated, heat-sensitive inhibition of autophagy by *Xcv*. Taken together, our detailed metabolic profiling showed a strong effect of the applied stress (combinations) on metabolites mainly involved in primary (energy) metabolism. For most sugars and urea, the impact of *Xcv* infection was effector-dependent and absent in *hrcV*. This effector-dependent effect was mimicked in *hrcV* by inhibition of autophagy.

## Discussion

The simultaneous occurrence of environmental stresses, such as heat, with pathogen infection poses a widespread threat to plant health. Understanding how plants navigate these dual challenges is crucial to enhancing crop breeding initiatives and creating robust varieties. In this study, we highlight autophagy as a fundamental mechanism that bestows stress-specific capabilities upon plants.

Response to combined abiotic-biotic stress is very complex, involving the interaction of two living organisms (the plant and the pathogen) as well as the additional level of abiotic stress (Kissoudis et al., 2014). Subjecting the plants to heat while concurrently exposing them to *Xcv* implies that both the plants and the bacteria are experiencing above-optimum temperatures. We tried to reduce the level of complexity by applying the combined heat-*Xcv* stress as a consecutive stress where the bacteria are not experiencing direct heat stress. Studies investigating combined stresses usually focus on a single time point; however, we believe that our experimental system of a consecutive stress setup reflects a natural situation more accurately where stress may also occur sequentially rather than simultaneously.

### Active autophagy in the host before infection facilitates bacterial growth

We observed that *Xcv* growth after infection was enhanced by preceding heat stress in the pathogen’s native host, tomato, as well as the model plant *N. benthamiana*, in an autophagy-dependent manner (Fig1A,B). Enhanced growth of another bacterial phytopathogen, *Pseudomonas Pst* DC300, at moderately elevated temperature during infection has been described before (Huot et al., 2017). As in our case, the increased growth depended on the delivery of bacterial effector proteins (Fig3C). We can now link this enhanced growth to bacterial manipulation of autophagy. Interestingly, over-expression of *ATG7* prior to infection yielded similar results, suggesting that other abiotic stresses inducing autophagy (such as drought or salt stress; (Avin-Wittenberg, 2019)) might elicit similar susceptibility to bacterial

The specific effector utilizing the pre-activation of autophagy by heat to facilitate bacterial growth is still unknown. So far, one *Xcv* effector, XopL, has been shown to regulate autophagy. Yet, it inhibits autophagy rather than utilizing it (Leong et al., 2022b). Our findings highlight the debate regarding the dual role of autophagy in pathogen resistance, promoting or inhibiting pathogen growth in a context-dependent manner (Mostowy, 2013). One possibility is that an effector other than XopL utilizes autophagy pre-activation to promote growth, which could not be identified in conditions where autophagy is not pre-activated. Another possibility is that the role of autophagy in immunity is indeed context-dependent, i.e., the autophagic response integrates several signals to produce a specific output.

### ATG8 confers specificity to combined plant stress responses

Autophagy can selectively remove damaged or unwanted cellular components. The selectivity is conferred by target binding to autophagy receptors, which, in turn, bind ATG8 (Stephani and Dagdas, 2020). Thus, although many conditions induce autophagy, the autophagic cargo and the resulting degradome are expected to be unique to each condition. Since plants have multiple ATG8 isoforms, it has been proposed that these isoforms play a crucial role in shaping the stress-specific response and selectivity of autophagy (Kellner et al., 2017, Lin et al., 2023, Zess et al., 2019). There are two main methods to study autophagy activation using GFP-ATG8. The GFP tag enables autophagosome visualization by confocal microscopy. In addition, the ratio between GFP-ATG8 and free GFP by Western blot allows us to estimate ongoing autophagic flux (Klionsky et al., 2021). We could not detect free GFP in our *N. benthamiana* Western blots, apart from plants exposed to dark conditions (Fig2A-C). In fact, we often and repeatedly only detected an unknown but GFP-specific degradation band. Similar bands were reported in other plant species (Zhou et al., 2023), yet their existence was not explained. This novel band is clearly connected to the degradation of GFP-*St*ATG8, and is absent without GFP-*St*ATG8 protein. Therefore, we decided to use this degradation band instead of free GFP to calculate the ratio between its intensity and GFP-*St*ATG8.

In our study, the seven *St*ATG8 isoforms exhibited stress-specific responses based on the type of stress (Fig2A-C). Carbon starvation triggered a uniform degradation of all isoforms, whereas *Xcv* infection or heat differed in ATG8 activation pattern. Bacterial infection is hampered by removing effector proteins via autophagy (Leong et al., 2022b). Accordingly, only selected *St*ATG8 isoforms were activated by *Xcv* infection. By contrast, under heat stress, *St*ATG8 isoforms differed in degradation timing. Heat stress generally causes protein aggregation, but proteins vary in their ability to withstand elevated temperatures. We suggest *St*ATG8 isoforms sensitivity and, thus, their need for clearance. One of the unique structural features of ATG8 proteins is an N-terminal helix domain (Bu et al., 2020). This domain determines the binding specificity to ATG8-interacting proteins, and its sequence is not conserved in plants (Zess et al., 2019). Thus, preferential binding of ATG8 isoforms to different targets and selective autophagy receptors seems only logical (Kellner et al., 2017). Besides differential activation of ATG8 proteins, expression of *AtATG8* genes differed when exposed to starvation or (a)biotic stress (Wang et al., 2019, Chiu et al., 2023, Leong et al., 2022b), further confirming our finding of stress-specific involvement of *St*ATG8 isoforms.

GFP-*St*ATG8-3.1 was involved in the response to heat stress and *Xcv* infection (Fig2B,C). Yet, when exposed to a combined heat-*Xcv* challenge, the activation did not mimic a combination of both stresses (Fig2E). Hence, the assumption that combined stresses equal the sum of the involved single stresses falls short of the complexity of natural systems. There is a possibility that combined stress activates another suite of *St*ATG8 isoforms than the single stresses. The Western blot results strengthen our infection assays with *Xcv*, suggesting that the autophagic response is tailored to integrate several stresses.

### Changes in metabolites as an underestimated effect of autophagy influencing bacterial performance

Nutrient availability plays a major role in pathogen growth. We thus asked whether previous heat stress affected nutrient availability for the invading pathogen. Due to technical constraints, our metabolic profiling focused on a sinlge time point, missing the dynamic regulation of metabolism. In addition, the large variation and lack of significant differences were disappointing. The experiment involved ten treatments and six biological replicates per treatment. The procedure was carefully controlled, including chamber transfers for temperature treatment and manual syringe infiltration for *Xcv* and *hrcV* treatments. However, potential changes in light quality and humidity between the experimental chamber and the cultivation room might have influenced metabolite levels. Additionally, the time-consuming tasks of harvesting two leaves per plant made precise 1-hour (60 min) sample collection impractical, resulting in a more realistic time range of 55 to 75 mins. However, we could come to several conclusions linking autophagy, bacterial effectors, and plant metabolism.

Nutrients in the apoplast available for bacterial nutrition are mainly sugars and amino acids (Rico and Preston, 2008, Fatima and Senthil-Kumar, 2015). Sugars and amino acids were generally reduced in the tested leaf tissue as a result of heat stress and also after infection. This observation is not surprising, given that stress responses are energetically costly (Signorelli et al., 2019). While we saw this reduction in amino acids upon *Xcv* infection (Fig4C), we cannot confirm it results from bacterial manipulation of autophagy or bacterial consumption of these metabolites. Urea, linked to amino acid catabolism (Witte, 2011), displayed an opposite response to stress. While amino acid levels decreased upon infection, heat stress alone had no major effect on urea levels, and *Xcv* infection or combined stress even slightly reduced urea levels (Fig5D). Interestingly, this reduction resulted from bacterial manipulation as levels increased significantly following *hrcV* infection. More importantly, this increase was absent when *hrcV* infection was combined with autophagy inhibition. Amino acids are recycled from proteins degraded via autophagy.

In addition, components targeted by bacterial effectors to manipulate metabolite recycling via autophagy might be heat-sensitive. When *hrcV* infection followed heat stress, urea levels were reduced, resembling *Xcv* infection. The absence of increased urea in combined *hrcV*-heat stress can also be explained on a more general immune response level. Salicylic acid (SA) is the dominant phytohormone involved in the plant’s defense against (hemi)biotrophic pathogens. This SA response is heat sensitive in Arabidopsis and tobacco (Wang et al., 2009). Loss of pathogen-induced SA was described as a central mechanism underlying enhanced bacterial growth of *Pseudomonas Pst* DC300 at elevated temperatures (Huot et al., 2017). Without an energy-intensive immune response, amino acids would not be channeled into energy production, failing to increase urea levels.

*Xcv* infection at least partially manipulates sugar levels in an autophagy-dependent manner, which seems to be mimicked by heat stress (Fig6). During infection, phytopathogens can adjust their metabolic machinery towards the most abundant nutrients in the host. *Xcv* was shown to modulate its manner of active nutrient acquisition (Fatima and Senthil-Kumar, 2015). So-called TBDTs (TonB-dependent transporters) are involved in uptake of carbohydrates, with genomes of several *Xanthomonas* species having a high copy number of TBDT genes (Blanvillain et al., 2007, Dugé de Bernonville et al., 2014, Pieretti et al., 2012). Moreover, pathogens like *Xcv*, colonizing the apoplast, release effector proteins to manipulate plant cells and optimize nutrient availability, e.g. by facilitating sugar efflux through induction of gene expression of sugar transporters (Chen et al., 2010, Eom et al., 2015, Zhou et al., 2015).

In addition to effectors influencing nutrient efflux, pathogens target autophagy directly (Üstün et al., 2016, Lal et al., 2020, Leong et al., 2022a). The reduction of sugar levels is partially dependent on effector proteins, as the *hrcV* mutant was unable to reduce sugar levels comparable to *Xcv*. Inhibition of autophagy by 3-MA led to *Xcv*-like sugar levels after *hrcV* infection, suggesting that the reduced sugar levels after Xcv infection resulted from autophagy inhibition by bacterial effectors. The autophagy-dependent manipulation of sugar levels is no longer observable when *hrcV* infection follows heat stress. Since heat stress affects autophagic activity (Zhou et al., 2014) and thereby also metabolite recycling, manipulation of autophagy might be obsolete under these conditions. Another interpretation is that, as mentioned above, the effector targets are heat sensitive, and this manipulation is no longer needed when the infection follows a heat stress. However, the effect of *Xcv* on sugar levels does not solely depend on autophagy, as autophagy inhibition by 3-MA did not alter the *Xcv* sugar phenotype. We thus postulate that manipulation of autophagy is not the only mechanism underlying pathogen success. This observation also explains why *Xcv* growth benefits from preceding heat stress whereas *hrcV* growth does not (Fig1B,C&3C). While the *hrcV* sugar phenotype can be compensated by preceding heat stress, the *hrcV* growth phenotype is unaffected, presumably because there are other effectors contributing to pathogen success than those targeting autophagy. As *hrcV* can not secrete any effectors to the plant, future studies using more specific effector mutants might be able to discern the particular effects of bacterial effectors in autophagy-dependent nutrient availability.

Our findings underscore the specificity inherent in the autophagic response, particularly regarding ATG8 isoforms. We demonstrate that combined stresses do not equal the sum of their counterparts, specifically focusing on autophagy at the nexus of simultaneous challenges. It will be interesting to see if the specificity of the autophagic response, especially in the case of stress combinations, also translates to the autophagic cargo level. The influence of heat stress on autophagy, seen in its facilitation of bacterial growth, establishes it as a key link between environmental factors and plant-microbial interactions. This connection underscores the broader ecological implications of autophagy in plant defense and unveils potential targets for enhancing stress resilience. As autophagy functions in many aspects of plant life, it is not possible to distinguish its roles in nutrient supply, abiotic stress, and pathogen defense. As our results show, it is possible that other, still unknown factors contribute to autophagy-dependent bacterial growth following heat stress.

## Acknowledgments

We thank the Rachel Green and Omri Finkel groups at HUJI for assistance with heatable phytochambers and Elena Minina, Johannes Stuttmann, Yasin Dagdas, and Guido Sessa for sharing transgenic seeds, bacterial strains and plasmids. Fig. 1A, 2D, 3A and 4A were created with Biorender (https://www.biorender.com). The project was funded by a Walter-Benjamin

## Author Contributions

HS and TAW designed the research; HS, EK, YS and TAW performed research experiments; HS, YS and TAW analyzed data; HS and TAW wrote the manuscript with input from all authors.

---

**Supplemental Figure S1:**
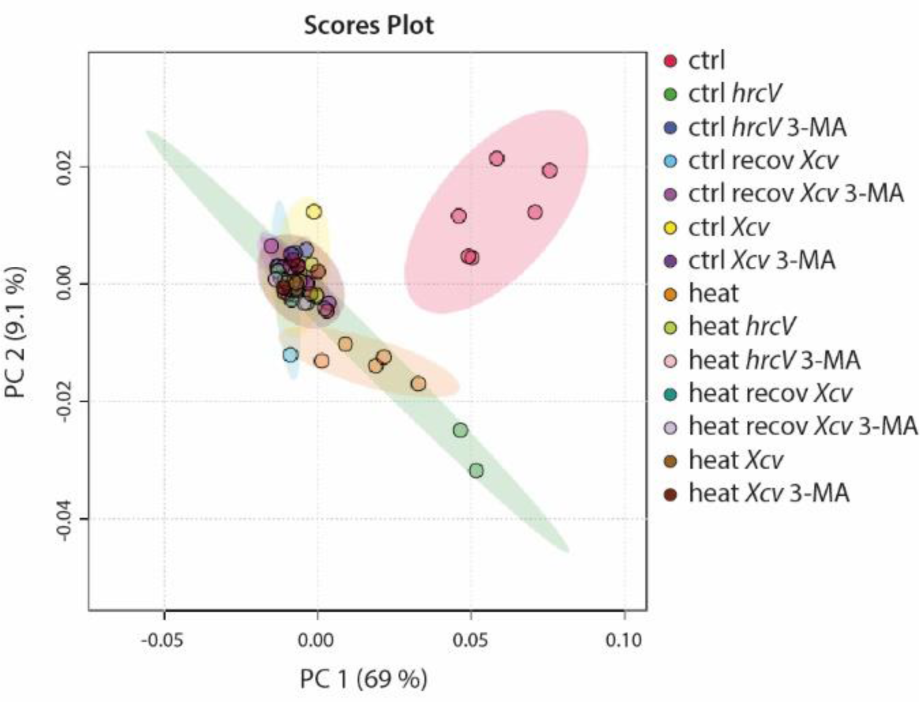
Principal component analysis of the metabolome as in Figure4B including heat recovery samples

**Supplemental Figure S2:**
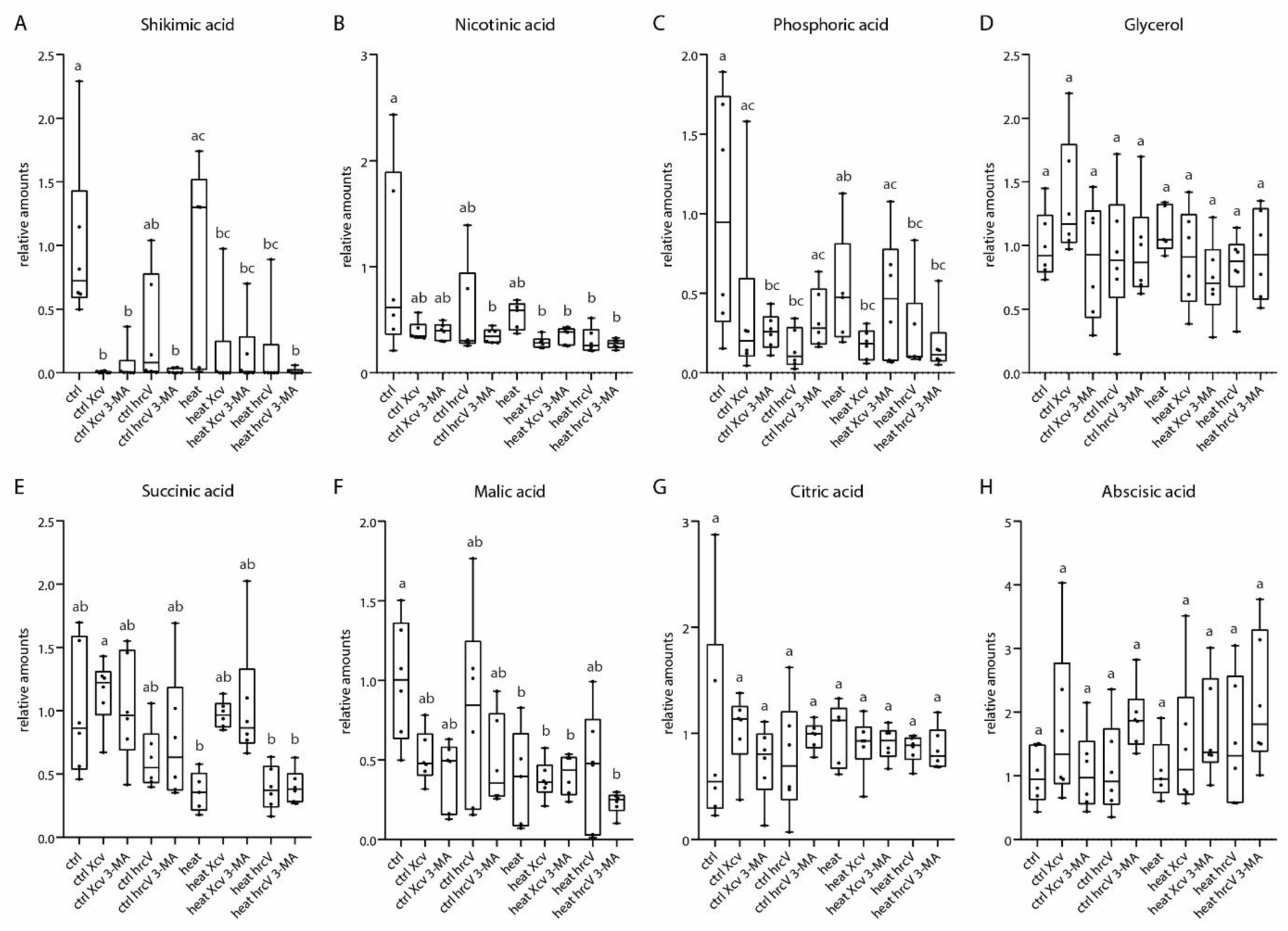
Detailed overview on effects of *Xcv* infection and heat on the levels of organic compounds. Boxplots showing levels of Shikimic acid (**A**), Nicotinic acid (**B**), Phosphoric acid (**C**), Glycerol (**D**), Succinic acid (**E**), Malic acid (**F**) Citric acid (**G**), and Abscisic acid (**H**) in response to *Xcv*, *hrcV*, and heat treatment, respectively (*n* = 5-6). Statistical analysis was performed by One-way ANOVA and Turkey’s multiple comparison test.

**Supplemental Figure S3:**
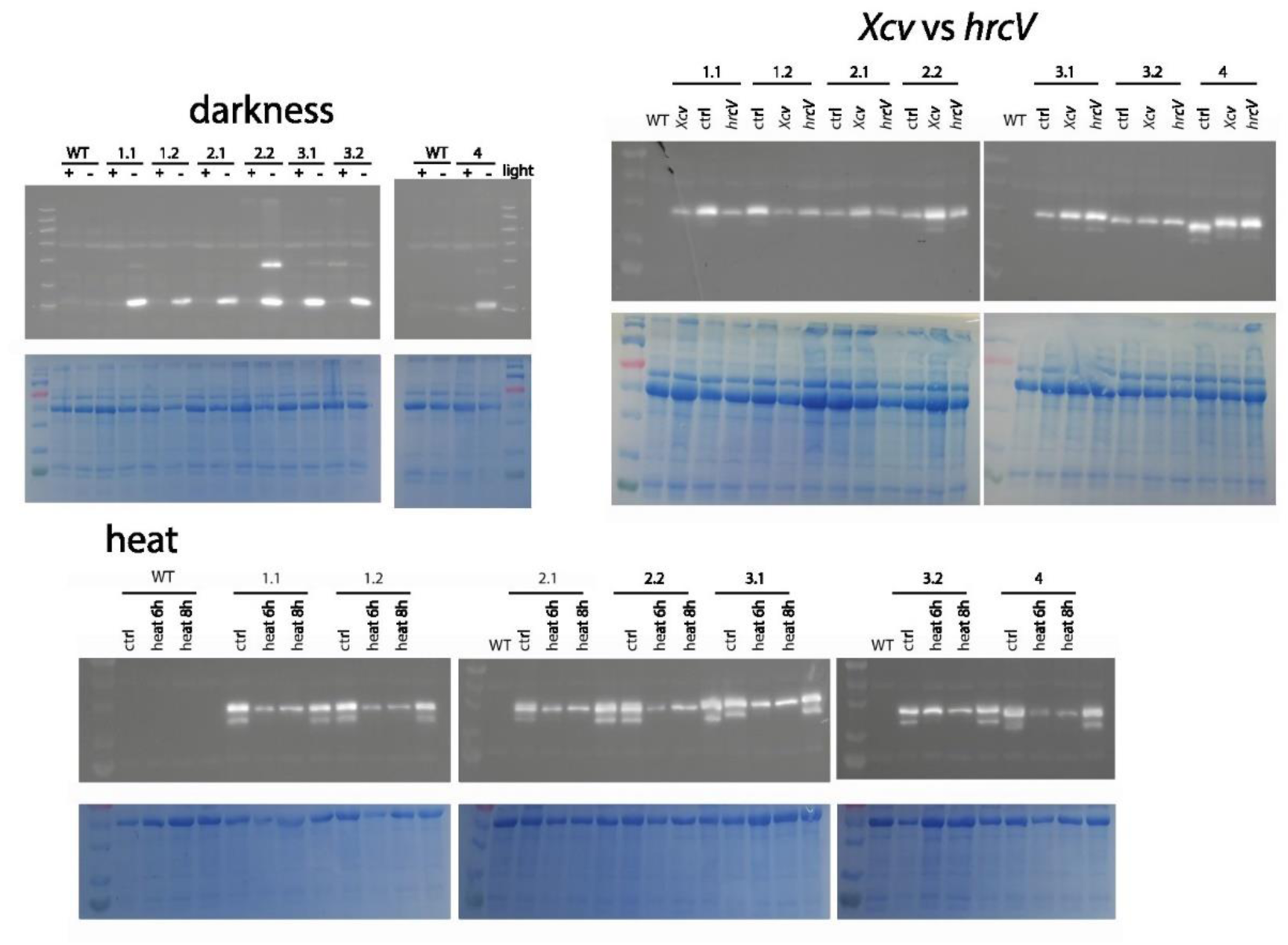
Original blots depicted in Figure 2

